# Genomic architecture of nestmate recognition cues in the desert ant

**DOI:** 10.1101/2023.11.08.566184

**Authors:** Pnina Cohen, Shani Inbar, Eyal Privman

**Author notes:** Equal contribution (Pnina Cohen, Shani Inbar).

## Abstract

Nestmate recognition is the basis for cooperation within social insect colonies. Quantitative variation in cuticle hydrocarbons (CHCs) is used for nestmate recognition in ants and other social insects. We carried out a Genome Wide Association Study (GWAS) of CHCs in the desert ant *Cataglyphis niger* by sampling 47 colonies, fully sequencing six workers from each colony, and measuring the relative amounts of their CHCs. Under the Gestalt colony odour model, social interactions between nestmates, in which CHCs are being transferred and mixed, are essential in creating a uniform colony CHC profile.

Therefore, we carried out a second GWAS between the colonies and their uniform Gestalt odour by averaging nestmate genotypes and comparing them to their averaged CHC amounts. Our results are in line with the Gestalt model. Together, the two analyses identified 99 candidate QTLs associated with 18 of the CHCs. Thirteen clusters of two to four QTLs located within 10cM from each other were identified, seven of which contained QTLs from both analyses. We conclude that nestmate recognition cues are a complex quantitative colony-level trait with a significant genetic component to their phenotypic variation and a highly polygenic architecture.

## Introduction

Colonial identity is the foundation and a prerequisite for any social interaction in insect societies. The ability to recognize nestmates and to discriminate them from aliens is needed for protecting the colony’s food reserves and brood from being plundered and for preserving territorial control over resources around the nest. In most social insect species, nestmates have high genetic relatedness, in keeps with Hamilton’s principles of kin selection and inclusive fitness^1,2^, and nestmate recognition is equivalent to kin recognition. A major class of chemical cues that facilitate nestmate recognition are lipids found on the cuticle, mainly n-alkanes, methyl-branched, and unsaturated hydrocarbons^3–8^. Insect cuticular hydrocarbons (CHCs) chains vary in length (up to 60, but mostly between 20 and 40 carbons), number and positions of methyl branches, and number and positions of double bonds^9^. Social insect species generally display relatively large numbers of different CHCs^10^. While different insect species have qualitatively distinct mixtures of CHCs, intraspecific variation of CHC mixtures is chiefly quantitative. The relative amounts of CHCs are the chemical profile that make up its identity^11–13^. In ants, the numbers of different CHCs are especially large, even in comparison to other social insects^10^, with dozens of distinct CHCs on the body surface of a single individual.

Intra-colony CHC composition variability is smaller than inter-colony variability^6,14^, and some studies reported that chemical distances (dissimilarity) between CHC profiles of colonies correlate with aggression levels between members of these colonies^15–17^. Diet, nest material, and other environmental factors were shown to influence ant CHC profiles^18–20^ and thereby contribute to nestmate recognition and inter-colony aggression. Evidence from various studies indicates that genetic factors are also important contributors to the colony-specific composition of CHCs (reviewed by Sprenger and Florian^21^). Correlation between genetic and chemical distances was reported for the yellow crazy ant^15^, the Argentine ant^17^, and the clonal raider ant^22^. Provost^23^ conducted controlled crosses of *Leptothorax lichtensteini* to produce unfamiliar ants with varying degrees of genetic relatedness, demonstrating aggression is correlated with genetic distance. A genetic component to nestmate recognition is also supported by cross-fostering experiments in *Formica rufinarbis*^24^. While these studies provided support for the hypothesis that CHCs are affected by genetic polymorphism, so far no study has identified specific genomic loci that may be responsible for such genetic effects.

Two models were suggested for the mechanism that creates a colony specific CHC profile^25^. In the individualistic model, all CHCs found on an individual are self-produced. Under this model, the similarities between nestmate CHC profiles may be explained by shared genetics between the closely related colony members and by their shared environment. The alternative model is that of the colony Gestalt odour, in which CHCs are transferred and mixed between colony members via allogrooming and/or trophallaxis (transfer of food)^5,24,26–28^ . This serves to reduce intra-colony variability, which may prevent nepotism in colonies with some kin structure (e.g. in patrilines of a monogyne, polyandrous colony). These two alternative hypotheses were examined in multiple studies, some of which support either of the two models. For example, when colonies of *Cataglyphis Iberia* ants were divided into two groups their CHC profiles diverged and inter-group aggression transpired^29^. The CHC profiles converged after the separated groups were reunited, as expected under the Gestalt model. In another study, on *Rhytidoponera confuse* ants, levels of aggression towards nestmates were measured after members of two colonies were caged together^30^. This study concluded that the individualistic component is a greater contributor to the colony chemical identity than the homogenizing Gestalt effect.

The uniform CHC profile of a colony is a colony-level phenotype, a phenotype which is the combined output (the average or any other function) of the individual phenotypes of nestmates. As with any other phenotype, colony-level phenotypes may be affected by internal factors such as genetics, epigenetics, animal size, and its health, and external factors such as nest material, diet, predation pressure, weather, and seasonality. Other examples of colony-level phenotypes in social insects include alternative social structures, such as the monogyne versus polygyne social structures in fire ants^31^, collective activities such as foraging and nest construction^32,33^, and social immunity, the defensive actions taken against pathogens and parasites^33–35^. In ants and other eusocial insects, a colony is typically made of close relatives that may be regarded as a single colonial organism or a superorganism^36,37^. Therefore, it is useful to consider a colony-level genotype as a high-ploidy genotype of a single entity. These collective genotypes may be studied for their association with any collective colony phenotype, in our case, the homogenized CHC profiles.

Here, we carried out the first Genome Wide Association Study (GWAS) to search for the genetic basis of among-colony variation in CHCs. Based on a pilot study on the power of alternative approaches for executing a colony-level GWAS^38^, we sampled 47 colonies of the ant *Cataglyphis niger* and fully sequenced the genomes of six workers from each colony. We tested for association between genomic polymorphism and the quantities of 34 CHCs using two approaches. The first was a standard GWAS of individual genotypes and phenotypes. We then conducted a colony-level GWAS, motivated by the Gestalt model. In the second analysis, we tested for associations between the colony-level genotypes, represented by the colonial allele frequency in each locus, and colony-level (averaged) CHC profiles. These analyses revealed candidate loci that underlie inter-colony variation in CHC profiles, which constitute the genomic basis for nestmate recognition cues.

## Materials and methods

### Study population

This study used a population sample from Betzet beach on the northern Israeli coastline. This population was previously referred to as *Cataglyphis drusus*^39^, but our recent species delimitation study suggested that this is the same species as *C. niger*, because these populations are not differentiated by their nuclear genomic DNA^40^. We note that some populations of *C. niger* include polygyne colonies (headed by multiple egg-laying queens)^40^ and that our recent study revealed that this social polymorphism is associated with a supergene (Lajmi *et al.* in prep). However, the study population used here is purely monogyne (each colony is headed by a single queen)^39,40^. The supergene is always homozygous (MM) in monogyne samples (and always heterozygous MP in polygyne samples), so we do not expect it to affect the population genetics and the performance of GWAS in this study population. *C. niger* queens are mated with 2-9 males (polyandrous), resulting in an average of 0.26 within-colony genetic relatedness^39,40^.

### Sampling scheme

We collected ants from nests dug along a 4 km transect in Betzet beach (from to N33.07868, E35.10705 to N33.05162, E35.10245; Figure 1) between April 24 and May 10, 2016. At least 30 workers were sampled from each of 50 nests, 47 of which were used in this study (N35.10297, E33.05339 - N33.05162, E35.10245). The other three nests were used in the pilot stage of this GWAS^38^.

**Figure 1.**
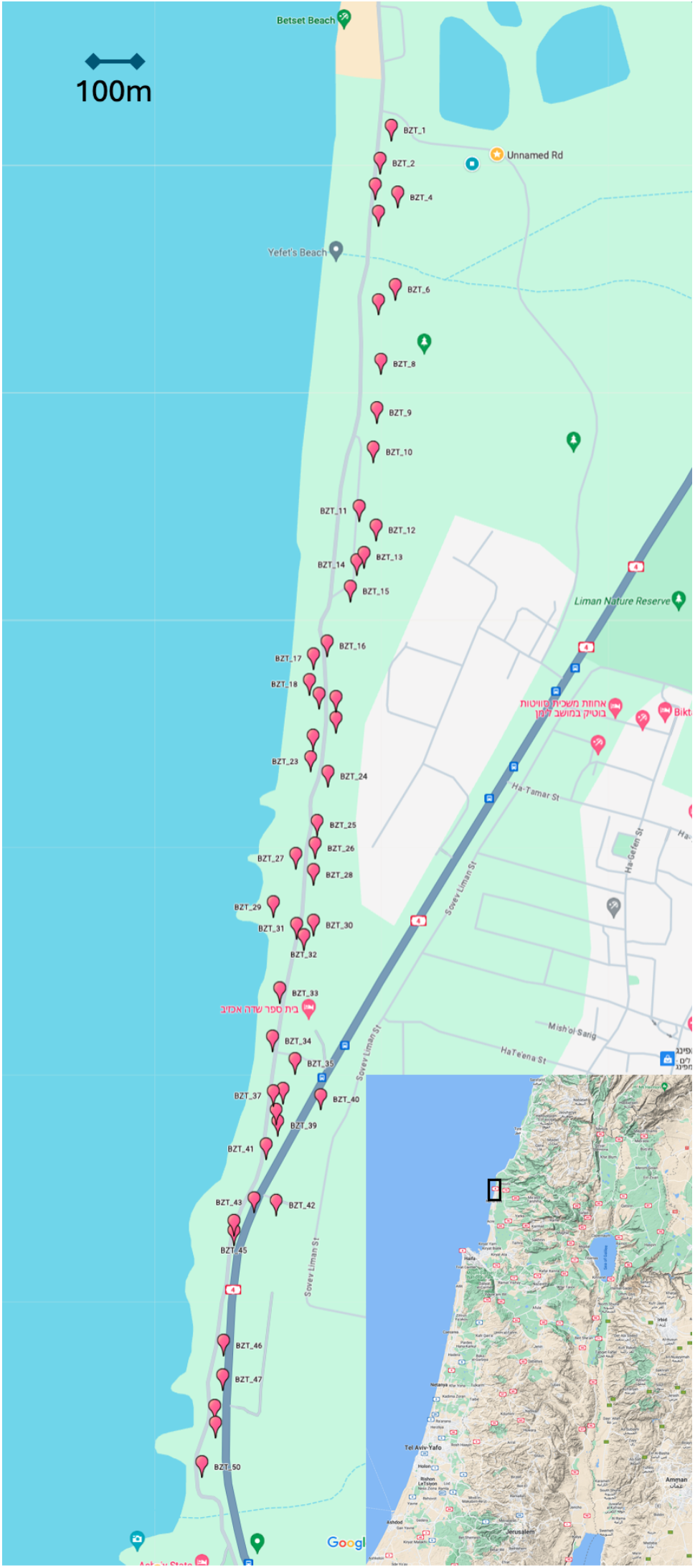
Map of 50 sampled nests along a transact in Betzet Beach in northern Israel (from to N33.07868 E35.10705 to N33.05162 E35.10245).

### Behavioural assays

There exists extensive literature on nestmate recognition studies in *C. niger* showing that CHCs are a major component of the chemical cues that workers use to discriminate non-nestmates from nestmates^6,28^. Behavioural assays were previously conducted in our study population, demonstrating that workers from different colonies discriminate nestmates from non-nestmates even between nests that are only 100m apart^40^. We conducted dyadic aggression assays to compare encounters between nestmates and between non-nestmates. The results confirm that the nests we sampled indeed exhibit the ability for nestmate recognition. See Inbar and Privman^41^ for the details of this behavioural assay.

### Pilot study, experimental design, and power analysis

Before sequencing and analysing our GWAS samples we conducted a pilot study to determine which experimental design gives maximal statistical power to detect QTLs for colony-level traits. In this pilot, we compared different combinations of population genomic methodologies and approaches, and also used to empirical results as a basis for a simulation study to assess statistical power. Please see the full details in Inbar *et al.*^38^. Based on the conclusions from this study we chose to conduct whole genome sequencing on six individual samples per colony. Naturally, the more samples the better, but the simulations showed diminishing returns for increasing the number of samples per colony. We note that this is true even though our study population consists of multiply mated queens (we simulated up to ten matings per queen, in line with estimates of 2-9 patrilines per colony in *C. niger*^40^). Based on these results we expect our GWAS to have 37% and 64% probability to detect QTLs that explain 23% and 52% of the variation in the studied trait, respectively (with a *p*-value of 5% after correction for multiple testing).

### Whole genome sequencing

For six individuals from each of the 47 nests, DNA was extracted from the head and abdomen (QIAGEN AllPrep DNA/RNA Mini kit) and the thorax was used for the chemical analysis of CHCs (see below). A uniquely barcoded genomic library was constructed for each sample following the standard protocol of the Illumina TruSeq Nano DNA Library Prep kit. Briefly, a Covaris S220 sonicator was used for DNA shearing aiming for 550bp inserts, ends were repaired, and the fragments were size-selected and purified using magnetic beads (supplied in kit). Then, an adenosine was added to the 3’ end of the blunt fragments, individually barcoded adaptors were ligated, and ligated fragments were amplified (eight PCR cycles). Libraries were pooled at equal quantities as three separate pools, each of which was sequenced in two lanes of a paired-end, 150bp reads, on a HiSeq X Ten Illumina sequencer (six lanes in total).

### Sequence data processing and genotype calling

#### Raw reads pre-processing

Whole-genome sequence data were successfully obtained for 276 of the 282 samples (four from colony 10, five from colonies 18, 19, 22 and 29 and six from each of the other colonies). A total of 5,941,275,998 reads of 150bp were sequenced, with an average of 21,526,362 reads per sample, which is an average genome coverage of 12X (0.5X – 28X). Duplicate read pairs were removed and low quality reads and adaptor contaminations were trimmed or removed using Trimmomatic^42^ version 0.36, leaving sequences that were at least 75bp long. The total number of remaining reads was 5,527,107,050 (93% of the raw data). Reads were aligned to the *C. niger* reference genome (version Cnig_gn2) using Bowtie2^43^ version 2.2.5 run with the ‘very-sensitive’ and ‘end-to-end’ settings. 3,216,552,155 reads (54% of raw reads) remained after the filtering out alignments with three or more mismatches or multiple mapping, which were defined as reads with a second best match that has less than twice the number of mismatches as the best match. HaplotypeCaller of the Genome Analysis Toolkit (GATK) package^44^ version 3.7 was used to create a Single Nucleotide Polymorphism (SNP) catalogue containing 2,792,460 SNPs.

In a first stage of filtering, we removed indels and SNPs with more than two alleles from the catalogue, leaving 2,413,918 SNPs. 16 samples (all four of colony 10, three of colony 11, two of colony 9 and one of colonies 2, 8, 15, 16, 25, 40 and 42) were removed because they were missing genotype data in more than 50% of the loci, leaving 260 individuals from 46 colonies in the analysis.

#### SNPs catalogue for GWAS

In a second stage of filtering we required for each locus to have genotype calls for at least 80% of the samples with a minimum number of 6 reads per genotype call, a maximum of 250, and a minimum minor allele frequency (MAF) of 15%. In order to remove false SNPs that are the result of collapsed repetitive sequences in the reference genome assembly, a control for these was made using three haploid male *C. niger* sampled in Betzet Beach (33°4′40.88″N/35°6′33.97″E) that were fully-sequenced with high coverage (177X, 55X and 72X). This was done by creating a blacklist of loci where one or more of these haploid samples appears to be heterozygous, which indicates an artifact because male ants are haploid. All SNPs within 600 bp of such heterozygous loci were removed from the catalogue. Additionally, excessively covered SNPs, defined as loci with an average number of reads per sample > 20 (average genome coverage + 2 x standard deviation – see Supplementary Figure S1A). We then removed 14 samples that had more than 60% missing genotypes after this filtering, and as a result had to exclude colony 9 because it had only two samples left. We were left with 244 individuals from 45 colonies (three from colony 11 and 29, four from colony 7, 8, 25 and 32, five from colonies 2, 14, 15, 16, 18, 19, 20, 22, 26, 30, 40 and 42 and six from all the rest) and 171,570 SNPs. The average coverage of this filtered SNP catalogue across the remaining 244 individuals was 10.2X (calculated by the actual number of reads covering the SNPs). Overall, 6,348,274 (15.2%) of all possible 41,863,080 locus X sample combinations did not have genotype call (missing data).

#### Colonial genotype processing and calculation

For the colony-level analysis, we required that every locus had genotype calls in at least three individuals in each colony, further reducing the number of SNPs to 171,355.

Overall, 659,721 (8.6%) of possible 7,710,975 locus X colony combinations did not have genotype call (missing data). We defined colony-level genotypes as the estimated allele frequency of the colony at every locus, which essentially means that a colony is treated as a single polyploid organism. For example, if the genotypes of six members of a colony are 00, 01, 01, 11, 11, 11, where 0 and 1 represent the two alternative alleles in a locus, their individual genotypes were coded in the input file for the GWAS analysis as 0, 1, 1, 2, 2, 2 and their colony-level genotype was calculated as an average:

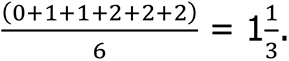

### Kinship inference

#### Individual-level kinship based on the whole-genome data

To create the kinship matrix on the individual level, we chose stringent filtering criteria to ensure low missing data rates and allowed low frequency alleles, which provide useful information for inference of identity-by-descent (IBD) of haplotypes. We filtered the SNP catalogue of the whole-genome sequenced samples (after first stage filtering) so that each locus had genotype calls in at least 88% of the samples with a minimum of 6 reads, a maximum of 250, and a minimum MAF of 1%. Here the MAF cutoff was reduced relative to the GWAS analysis above because rare alleles are a powerful signal for kinship. A MAF of 1% still requires a minimum of two samples that share the minor allele (if homozygous; more if heterozygous), which makes it unlikely to be an artifact due to sequencing errors. We controlled for and filtered out reads that were mapped to suspected collapsed repetitive sequences in the reference as described above, and similarly removed 16 samples from the analysis due to insufficient genotype calls. This resulted in a catalogue of 88,238 SNPs. Using the program Beagle^45^, we imputed and phased the catalogue to the total of 9,634 phased segments. The refined-IBD algorithm^46^ was used to identify shared haplotypes between the samples. A kinship matrix was created based on the total length of IBD haplotypes shared between every pair of samples.

#### Colony-level kinship based on RAD-sequencing

To better estimate kinship between colonies, one individual from each of the 47 colonies was chosen to be sequenced with greater sequencing depth using a reduced-representation genomic sequencing approach: the double-digest Restriction-site Associated DNA sequencing (ddRAD-seq) protocol. The rationale for this is that kinship inference by detection of haplotypes that are identical by descent (IBD) between pairs of samples depends on low genotype error rates. Using ddRAD-seq we could sequence a smaller number of loci to a much greater depth (65X – see below), which reduces the genotype error rate.

#### Restriction-site associated DNA (RAD) sequencing

For one worker per nest, a library was constructed according to a ddRAD-seq protocol based on protocols from Peterson *et al.*^47^ and Brelsford *et al.*^48^. Briefly, DNA was digested by two different restriction enzymes (EcoRI as the rare-cutter; MseI as the frequent-cutter) and ligated to barcoded adaptors for multiplexing. Products were amplified using Q5 Hot Start Polymerase (NEB), with the number of cycles reduced to 20, the number of replicates increased to four, and starting DNA volume increased to 6ul. Also, primers and dNTPs were added to the final thermal cycle in order to minimize single-stranded or heteroduplex PCR products. The libraries were sequenced in one lane of a paired-end, 150bp reads, on a HiSeq X Ten Illumina sequencer.

#### Pre-processing of RAD-seq data

A total of 823,087,220 reads were sequenced from 47 workers (one from each of the colonies), which is an average expected depth of 65X (14X – 105X) based on the number of restriction sites in the reference genome. The raw reads were initially processed using the *Stacks* pipeline^49^ version 2.2. Low quality reads were discarded if their quality score dropped below an averaged phred score of 10 in a sliding window of 15% of the reads’ length. After this initial processing, 736,899,655 reads that were assigned to a sample remain (90% of raw data). The reads were mapped to the reference genome of *C. niger* (version Cnig_gn2) using *Bowtie2* and filtered using the same criteria as for the whole-genome data above. After this filtering, 718,157,408 alignments remained. We continued to analyse the mapped sequences using the *gstacks* program of the *Stacks* pipeline and created a catalogue containing 90,006 SNPs for the 47 samples.

#### Combining the catalogue with additional samples from across Israel

To increase the number of samples and thereby improve SNP calling, imputation and phasing, we made use of another data set that included 103 RAD-sequenced *C. niger* and *C. israelensis* individuals sampled across Israel (described in Rainer-Brodetzki *et al.*^40^). The combined catalogue that was created from the 150 individuals contained 346,940 SNPs.

The following filtering parameters were used: each locus was required to have a genotype call in at least 90% of the samples, with the minimal number of reads of 3, a maximum of 4000, and had a minimum MAF of 1.5%. We controlled for collapsed repetitive elements by using the blacklist described above and by removing excessively covered SNPs, which were defined as loci with a mean number of reads > 250 (mean coverage + 2 x standard deviation – see Supplementary Figure S1B). After filtering, 21,179 polymorphic sites remained in the combined catalogue.

#### Colony-level kinship inference based on RAD-seq data

As with the whole-genome data, the RAD-seq SNP catalogue was imputed and phased using Beagle and refined-IBD to a total of 502 phased segments across the genome.

Shared IBD haplotypes between individuals were identified and a kinship matrix was created based on the total length of IBD haplotypes shared between every pair of samples representing 47 colonies.

### Chemical analysis of cuticular hydrocarbons

#### Cuticular hydrocarbon (CHC) analysis

CHCs were measured for the same six workers from each of the 47 nests that were used for genomic sequencing. Thoraces were individually immersed in hexane to extract non-polar cuticular lipids from each sample separately. Initial analysis was conducted by gas chromatography/mass spectrometry (GC/MS), using a VF-5ms capillary column, temperature-programmed from 60◦C to 300◦C (with 1 min initial hold) at a rate of 10◦C per min, with a final hold of 15 min. Compounds were identified according to their fragmentation pattern and respective retention time compared to authentic standards.

Subsequently, all samples were assayed quantitatively by flame ionization gas chromatography (GC/FID), using the above running conditions. Peak integration was performed using the program Varian Galaxie (version 1.9) and relative amounts of each compound were calculated. The relative amounts of each CHC were calculated for each sample. We confirmed that different nests have distinct CHC profiles using a linear discriminant analysis (LDA p-value<0.0001; Supplementary figure S2). For the colony-level analysis, the relative amounts were averaged between the members of each colony.

#### Normalization of the CHC relative amounts

Outliers to the distribution of each CHC in each of the two datasets (individual relative amounts and averaged colony relative amounts) were removed. These were defined as the amounts that were 1.5 times higher or 1.5 times lower than the interquartile range (the difference between the 75th and 25th percentiles of the distribution). For each dataset, we normalized the distribution of the remaining values using the R package *bestNormalize*^50^, which chooses the best fitting normalization function for the dataset on the basis of the Pearson’s P statistic. See Supplementary Table S1 for more details.

#### CHC GWAS

In order to test for a genetic basis of variation in CHCs, we conducted a Genome Wide Association Study (GWAS) over the relative amounts of each CHC, in two levels. The first is the individual level, in which we used the catalogue of genotypes created for the 244 samples in 45 colonies, as described above, and their relative amounts of CHC. The second level is the colony level, in which we used the catalogue of colony-level genotypes created for the 45 colonies, as described above, and their averaged colony-level CHC relative amounts.

The standard practice for GWAS is the use of a Mixed Linear Model (MLM), in which the population structure and the genetic variation are represented as fixed effects, and kinship between samples as random effects. Here, for each dataset (individual or colony level), we to accounted for sample relatedness by the respective kinship matrix (whole-genome or RAD sequencing data for the individual- or colony-level datasets, respectively) and for population structure by a Principle Component Analysis (PCA) of the SNPs over 45 dimensions in the individual-level analysis and 8 dimensions in the colony-level analysis. The number of dimensions was chosen based on the fit of quantile-quantile (Q-Q) plots – see below.

#### Execution of GAPIT

Genomic Association and Prediction Integrated Tool (GAPIT)^51^ is an R package which executes the Efficient Mixed Model Association (EMMA)^52^ method, a computationally efficient algorithm for the study of genomic association using MLM. We ran the program for each CHC in each dataset separately using the Multiple Loci Mixed Linear Model (MLMM), which is implemented in GAPIT by a forward-backward stepwise MLM regression. In each step, the loci that were found to be significantly associated with the trait in the former step are added to the model as cofactors. Accounting for the effect of multiple loci on the trait reduces residual variation in the trait and allows for a greater statistical power in QTL detection. We evaluated how well the model accounts for population structure and kinship using Q-Q plots: plotting the negative logarithm of the assessed *p*-values and comparing them to the expected distribution of *p*-values under the null hypothesis of no association, as calculated by GAPIT. See Supplementary Figure S3. Of the 171,570 / 171,355 loci analysed, 155,815 / 155,807 were mapped to a linkage group on the genetic map^52^.

#### Empirical permutation test

To account for additional unknown confounding factors and other potential problems in the model that might lead to false inference of association, we performed a permutation test to empirically calculate a null distribution of *p*-values when no association between the genotype and phenotype exists: for each CHC, we randomly shuffled the trait values between the samples. Using the shuffled traits and leaving the other model parameters unchanged, we rerun GAPIT 100 times. We did this for both the individual-level and colony-level analyses. For each trait, the original *p*-values of the original run were re-evaluated by comparing them to the distribution of the *p*-values in the shuffled datasets. The *p*-values were further corrected for multiple testing using the Benjamini-Hochberg false discovery rate (FDR) method^53^. We report associations above multiple significance thresholds, with FDR ranging from 5% to 30%. While it is not common to report results with 30% FDR, we find it useful to include this lower threshold. For example, in the individual-level GWAS we report 69 loci above this threshold. While 30% of them are expected to be false positives, these results show that approximately 48 loci (70%) are associated with CHCs across the genome. We ask our readers to keep this in mind while considering our results.

## Results

We conducted a GWAS to identify the genetic basis of variation in CHCs. We tested for associations between the relative amounts of 34 CHCs and genotype calls at 171,570 SNPs identified through the whole-genome sequencing of 244 workers from 45 colonies of *C. niger* (3 – 6 ants from each colony). As part of the analysis, relatedness between samples is accounted for by a kinship matrix and population structure is accounted for using a PCA. This analysis was conducted both at the individual level and at the colony level, where the colony phenotype was defined as the averaged CHC level in the colony and the colony genotype as the allele frequency in the colony. Each locus was assigned with a *p*-value for its association with each of the traits, which we corrected for multiple testing into *q*-values.

### Population structure and kinship between colonies

It is important to account for population structure and kinship between samples before conducting a GWAS. We analysed population structure using principle component analysis (PCA) of RAD-seq data from one sample of each of the 47 colonies. This analysis revealed limited spatial variation along the north-south axis (Supplementary Figure S4). This may be due to limited dispersal during mating flights. Kinship among samples was inferred from the whole-genome sequencing data based on the total length of shared IBD haplotypes (Supplementary Figure S5A). As expected, and with very few exceptions, stronger kinship was identified between sample collected from the same nest (intra-colony kinship) relative to kinship between nests (inter-colony kinship). The inter-colony kinship between members of colonies 33 and 47 was found to be similar to intra-colony kinship. A possible explanation is that the queens of these two colonies were closely related, either a mother and a daughter (50% related) or full sisters (75% related). There is also high relatedness between some members of colonies 40, 41 and 42. And there appears to be some kinship among colonies 1 – 20 (northern part of the transect) and among colonies 32 – 47, in line with the limited structure observed in the PCA results. Kinship was also inferred from more deeply sequenced RAD-seq data from one representative of each colony. High sequencing depth is expected to give lower genotype error rate, which is important for kinship inference. Various levels of inter-colony kinship were observed, mostly between neighbouring nests, which also supports the hypothesis of short range dispersal (between colonies 16 and 19; colonies 34 and 37; colonies 35, 40, 41 and 42; and colonies 46 and 47; see Supplementary Figure S5B).

### Individual-level GWAS

The individual-level GWAS identified 13 QTLs that were found to be associated with one of the CHC with the corrected *q*-value < 0.1, three of which were mapped to genes (Figure 2 and Table 1). Two of the QTLs found associated with 5Me-C25 were only 4.6cM apart of on chromosome 15. Additionally, 3, 13, 39, and 69 loci were found to be associated with one of the traits with a *q*-value < 5%, 10%, 20%, and 30%, respectively. Of these, 22 were located within a gene or within 2000 bp upstream from a gene’s start codon (gene hits). We note that for the lower significance threshold 30% of the associations are expected to be false positives, so approximately 48 out of 69 putative QTLs are expected to be true positives. We also used Q-Q plots to assess the distribution of *p*-values under the null hypothesis of no association. These plots show that the association test was well-controlled for potential artefacts such as kinship or non-normal trait distributions that might result in false positives (Supplementary Figure S3). The associated CHCs are shown in the order of their appearance on the chromosome (individual-level QTLs in blue and colony-level QTLs in black)

**Figure 2.**
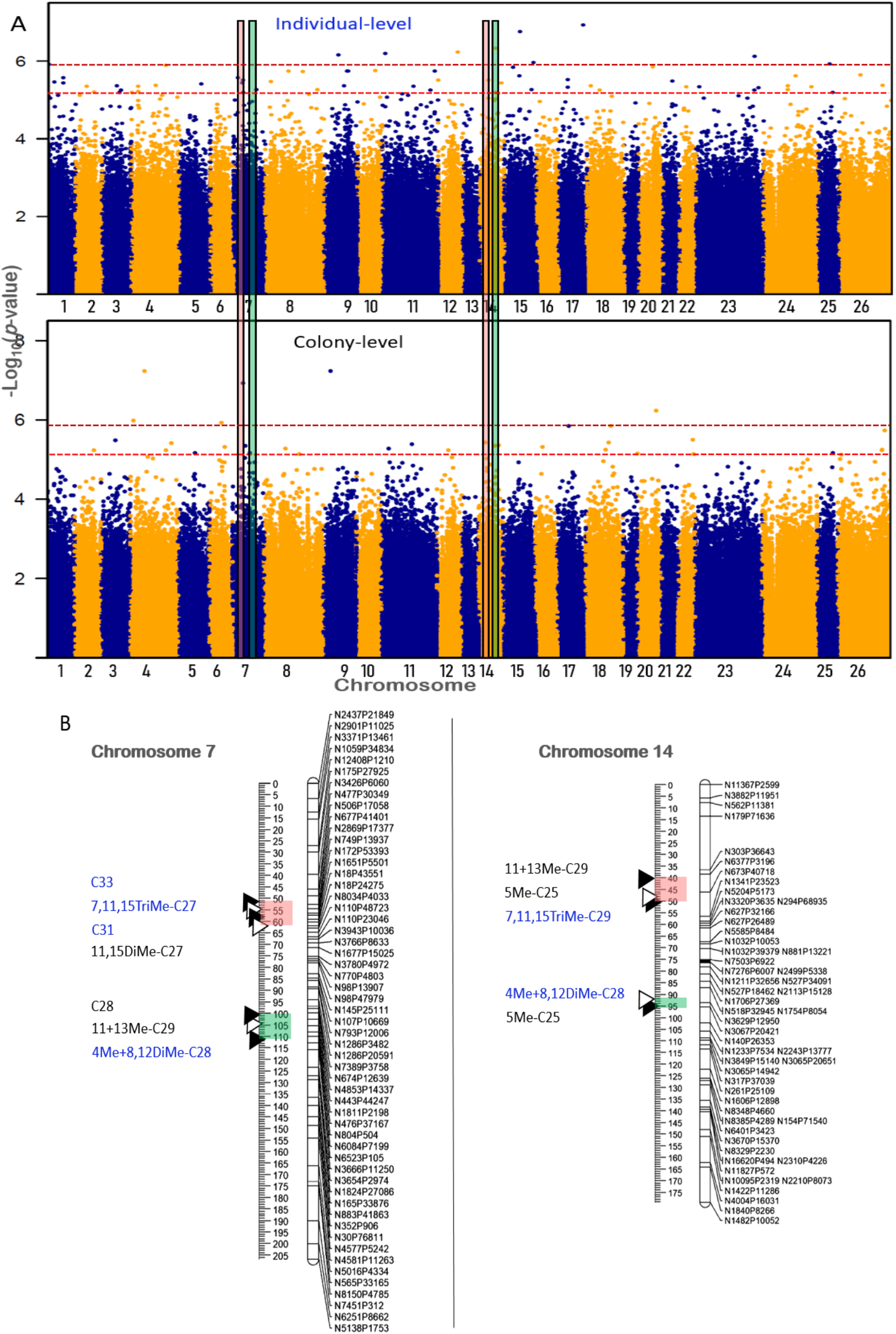
Manhattan plots of associations between loci and CHCs from the individual-level (A) and colony-level (B) GWAS. Loci above the maroon dotted line are statistically significant with false discovery rate (FDR) of 10%. Loci above the red dotted line are statistically significant with FDR of 30%. Four QTL clusters with QTLs identified in the individual-level and in the colony-level analyses are marked on chromosomes 7 and 14 and presented in detail (B).

**Table 1.**
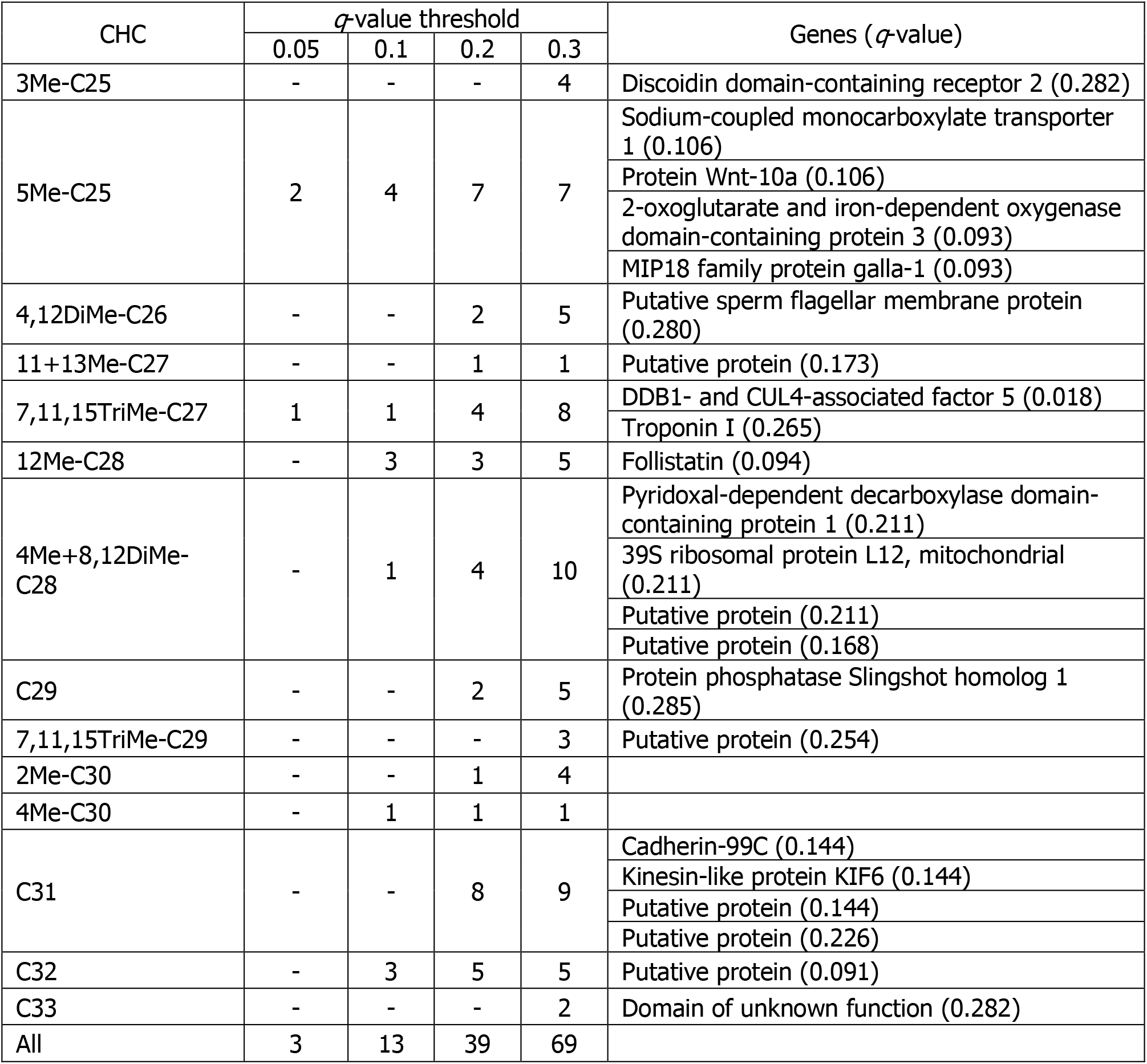
Candidate loci and genes that were found to be associated with CHCs in the individual- level GWAS

| CHC | *q*-value threshold |  |  |  | Genes (*q*-value) |
| --- | --- | --- | --- | --- | --- |
|  | 0.05 | 0.1 | 0.2 | 0.3 |  |
| 3Me-C25 | - | - | - | 4 | Discoidin domain-containing receptor 2 (0.282) |
| 5Me-C25 | 2 | 4 | 7 | 7 | Sodium-coupled monocarboxylate transporter 1 (0.106) |
|  |  |  |  |  | Protein Wnt-10a (0.106) |
|  |  |  |  |  | 2-oxoglutarate and iron-dependent oxygenase domain-containing protein 3 (0.093) |
|  |  |  |  |  | MIP18 family protein galla-1 (0.093) |
| 4,12DiMe-C26 | - | - | 2 | 5 | Putative sperm flagellar membrane protein (0.280) |
| 11+13Me-C27 | - | - | 1 | 1 | Putative protein (0.173) |
| 7,11,15TriMe-C27 | 1 | 1 | 4 | 8 | DDB1- and CUL4-associated factor 5 (0.018) |
|  |  |  |  |  | Troponin I (0.265) |
| 12Me-C28 | - | 3 | 3 | 5 | Follistatin (0.094) |
| 4Me+8,12DiMe-C28 | - | 1 | 4 | 10 | Pyridoxal-dependent decarboxylase domain-containing protein 1 (0.211) |
|  |  |  |  |  | 39S ribosomal protein L12, mitochondrial (0.211) |
|  |  |  |  |  | Putative protein (0.211) |
|  |  |  |  |  | Putative protein (0.168) |
| C29 | - | - | 2 | 5 | Protein phosphatase Slingshot homolog 1 (0.285) |
| 7,11,15TriMe-C29 | - | - | - | 3 | Putative protein (0.254) |
| 2Me-C30 | - | - | 1 | 4 |  |
| 4Me-C30 | - | 1 | 1 | 1 |  |
| C31 | - | - | 8 | 9 | Cadherin-99C (0.144) |
|  |  |  |  |  | Kinesin-like protein KIF6 (0.144) |
|  |  |  |  |  | Putative protein (0.144) |
|  |  |  |  |  | Putative protein (0.226) |
| C32 | - | 3 | 5 | 5 | Putative protein (0.091) |
| C33 | - | - | - | 2 | Domain of unknown function (0.282) |
| All | 3 | 13 | 39 | 69 |  |

### Colony-level GWAS

The colony-level GWAS identified three QTLs that were found to be associated with 5Me- C25, two of which with *q*-value < 0.05 (Figure 2 and Table 2). These QTLs were located on different chromosomes from the loci identified in the individuals-level analysis. Two additional QTLs were found to be associated with two other CHCs. There are 4, 5, 16, and 30 QTLs that with a *q*-value < 5%, 10%, 20%, and 30% that were found to be associated with CHCs, of these, six were gene hits. We note that for the lowest significance threshold 30% of the associations are expected to be false positives, so approximately 21 out of 30 putative QTLs are expected to be true positives.

**Table 2.** Candidate loci and genes that were found to be associated with CHCs in the colony-level GWAS.

| CHC | *q*-value threshold |  |  |  | Genes (*q*-value) |
| --- | --- | --- | --- | --- | --- |
|  | 0.05 | 0.1 | 0.2 | 0.3 |  |
| 5Me-C25 | 2 | 3 | 6 | 7 | Sterile alpha and TIR motif-containing protein 1 (0.142) |
| 11+13Me-C27 | 1 | 1 | 1 | 3 | Aminopeptidase N (0.276) |
| 11,15DiMe-C27 | 1 | 1 | 1 | 4 | Protein TANC2 (0.298) |
| C28 | - | - | - | 7 | Progesterin and adipoQ receptor family member 4 (0.227) |
|  |  |  |  |  | Protein gooseberry-neuro (0.261) |
| 11+13Me-C29 | - | - | 7 | 8 | Claudin tight junction (0.233) |
| 3Me-C29 | - | - | 1 | 1 |  |
| All | 4 | 5 | 16 | 30 |  |

Put together, 99 QTLs were found to be associated with CHCs with *q*-value < 0.3. The QTLs distribution across the genome was found to be significantly different than a uniform distribution in a Kolmogorov-Smirnov test (D = 5038.9, *p*-value < 2.2x10^-16^). We identified 13 clusters of two to four loci that were within 10cM long from each other. Seven of the clusters included QTLs from both individual- and colony-level analyses (Supplementary Table S2). Overall, 32 of the 99 QTLs were in clusters. There were additional three clusters, each containing 2 QTLs that were more than 10cM but less than 12cM from each other.

Of the gene hits, some are implicated in metabolism of amino acids. For example, pyridoxal-dependent decarboxylase domain-containing protein 1 contains a QTL for 4Me+8,12DiMe-C28. Other QTL genes are regulatory factors such as protein phosphatase Slingshot homolog 1 for C29. Other genes code for claudin, cadherin and kinesin, which may affect cell structures and functions.

## Discussion

We conducted a GWAS of CHC profiles in the desert ant *C. niger* on both the individual and the colony level. We detected a total of 99 loci associated with CHCs across the genome. These results show that the genomic architecture of nestmate recognition cues is highly polygenic. We found one or more putative QTLs for 18 of the 34 CHCs (*q*-value < 0.3): 14 in the individual-level analysis and 6 in the colony-level analysis. Two CHCs appeared in the results of both analyses (5Me-C25 and 11+13Me-C27). For the remaining 16 CHCs, either their QTLs were missed due to problems in the analysis itself, such as chemical measurement or sequencing errors, wrong model choice, inaccurate kinship inference, etc., or because the contribution of genetics to their quantitative variations was too low to be detected with limited statistical power of GWAS using our limited sample size. Nevertheless, this is the first report of QTLs for CHCs in any ant species.

While neither the individual- nor the colony-level analyses identified the same locus as being associated with more than one CHC, oftentimes QTLs of different CHCs were found close by in the genome. It has been shown that some chromosomal regions form distinct compartments in regards to their functionality and co-regulation^54,55^. We hypothesize that the genomic clusters identified may be topological or functional domains that are related to the synthesizing of CHCs. Insects synthesize hydrocarbons by elongating fatty acyl-CoA precursors to produce long chain fatty acids that are then converted to hydrocarbons^9^. The elongation reaction is responsible for regulating hydrocarbons lengths^56^ and thereby in determining the variation in quantities of various CHCs with a range of chain lengths. Another source of variation is the addition of methyl branches in different positions during the synthesis of the fatty acids. The candidate QTLs may be related to these biosenthetic mechanisms, perhaps as polymorphic regulatory factors. Further study of the candidate loci is needed in order to examine their relation to CHC synthesis.

The ability to discriminate between nestmates and non-nestmates depends on the differences between their CHC profiles^6,14–17^. We may expect that the more distinct their colony CHC profile is, the easier it would be for nestmates to identify each other.

Colonies with rare genotypes in loci that regulate CHCs would therefore be favoured by natural selection over colonies with common genotypes. Rare alleles would be selected for so long they are rare, thus maintaining and increasing the polymorphism in the population. Therefore, we may predict balancing (frequency-dependent) selection on genes that regulate CHCs, in a similar fashion as in other recognition systems for which polymorphism is advantageous for the discrimination of self vs. non-self, such as the MHC immunity system in mammals^57,58^ and the self-incompatibility system of plants^59^. This leads us to predict that CHC variation will increase in evolution through the generation of polymorphic genetic loci that affect CHC synthesis and/or their secretion on the cuticle, i.e. CHC QTLs.

Despite this prediction for the evolution of genetic polymorphism in nestmate recognition cues, environmental factors may have greater contribution than genetic factors to the phenotypic variation. In our analysis, only seven QTLs for four of the 34 CHCs were detected with a *q*-value < 0.05; two CHCs in the individual- and three in the colony-level analysis (QTLs for CHC 5Me-C25 were detected in both analyses). Diet was shown to influence the cuticular chemical profiles in the polygyne and polydomous colonies of *Formica aquilonia* ants, where nestmates are not necessarily related^19^. In comparison, the colonies of *C. niger* used in this study are monogyne^40^, and so nestmates are always related (at least 25% relatedness). Environmental odours also appear to determine aggression levels between lab-reared and field-collected workers of monogyne *Solenopsis invicta* colonies^20^ and similarly between lab-reared and field-collected polygyne *Rhytidoponera confusa* workers^18^. Considering the limited results from our GWAS, it appears that genetic factors have a lesser role than environmental factors in our study system.

Nevertheless, our analyses suggest that polymorphism in multiple genes contributes to CHC variation. The statistical power obtained from the sample size of 244 ants from 45 colonies limited our confidence in specific QTLs, yet we can conclude from the results that tens of QTLs affect CHCs. This observation is in line with studies in model insect species such as *Drosophila*, where larger sample sizes and more powerful experimental approaches are available. For example, a GWAS of CHCs conducted using 169 inbred lines from the *D. melanogaster* Genetic Reference Panel (DGRP) identified QTLs in 305 and 173 genes in females and males, respectively^60^. Also in *Drosophila*, it is likely that the sample size limits the statistical power of GWAS. The numbers of QTLs reported are likely underestimates because QTLs with smaller effect size are more difficult to detect. Therefore, we may conclude that CHCs have a complex polygenic genomic architecture, which is in line with their being complex quantitative traits.

The Gestalt model suggests that social interaction between colony members has an important role in determining colony identity. Social interactions have substantial impacts on many other phenotypes in insect societies. For example, when *Camponotus floridanus* workers were fed with juvenile hormone, an important regulator of growth and development in insects, the larvae they fed through trophallaxis were twice as likely to become larger workers^61^. In termites, social interactions of allogrooming and trophallaxis appear to contribute in determining the caste of developing brood^62^. The physical transfer and mixing of CHCs between nestmates may be a major factor shaping their chemical identity. In our colony-level analysis, transfer of CHCs between nestmates is accounted for by the averaging out of nestmate genetics and their CHC phenotypes.

This analysis resulted in fewer QTLs identified in comparison to the individual-level analysis, but in general with greater statistical significance, indicating more confidence in the colony-level model. This result may be interpreted as supporting the Gestalt model. Alternatively, it may be that the averaging of colonies’ genotypes and phenotypes helps in eliminating noise introduced in the chemical analysis and sequencing/genotype errors, and so results in a more robust outcome. Additional studies including behavioural analysis and manipulation may provide further insight into the significance of the Gestalt model and determine the exact nature of social effects on CHC profiles. Whichever is the case, our colony-level analysis was successful in revealing putative QTLs, indicating that this approach is appropriate for CHCs. Whether the Gestalt effect is present or not, genomic mapping of CHCs should be done at the colony level due to the shared genetics and environment of nestmates.

## Acknowledgements

We thank Abraham Hefetz for advice and access to equipment for chemical analysis of CHCs. We thank Jessica Purcell and Alan Brelsford for advice on RAD-seq. This study was funded by Israel Science Foundation grant number 646/15.

## Data Accessibility

Genomic sequencing data will be made public upon acceptance of the paper in the NCBI SRA database. Chemical data will be made public in the Dryad repository.

---

**Figure S1.**
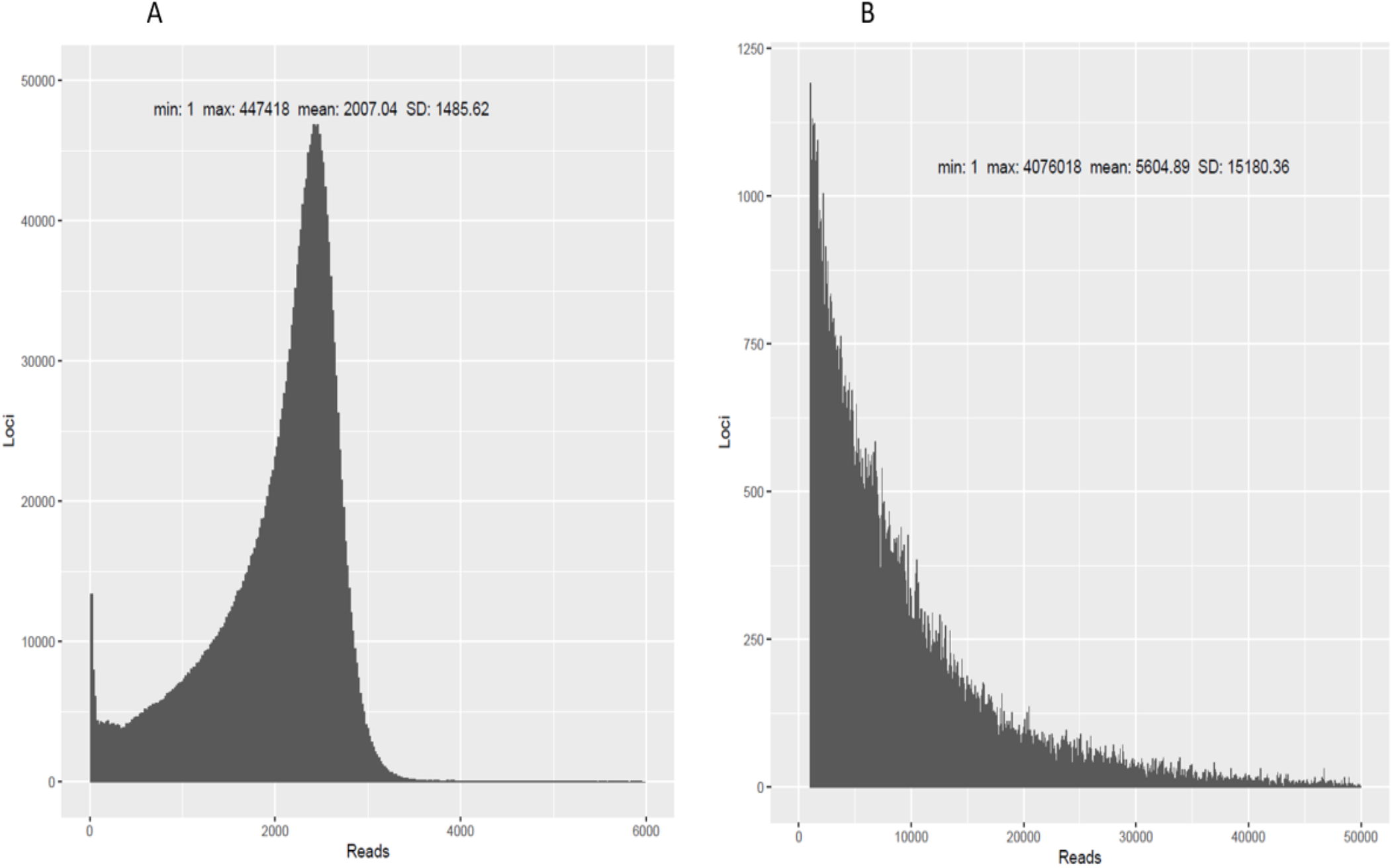
Distribution and summary statistics of the number of reads across unfiltered loci. (A) 276 fully sequenced samples from 47 colonies sampled in Betzet beach in northern Israel. (B) 47 RAD-sequenced samples of 47 colonies from which the fully sequenced samples also originate.

**Figure S2.**
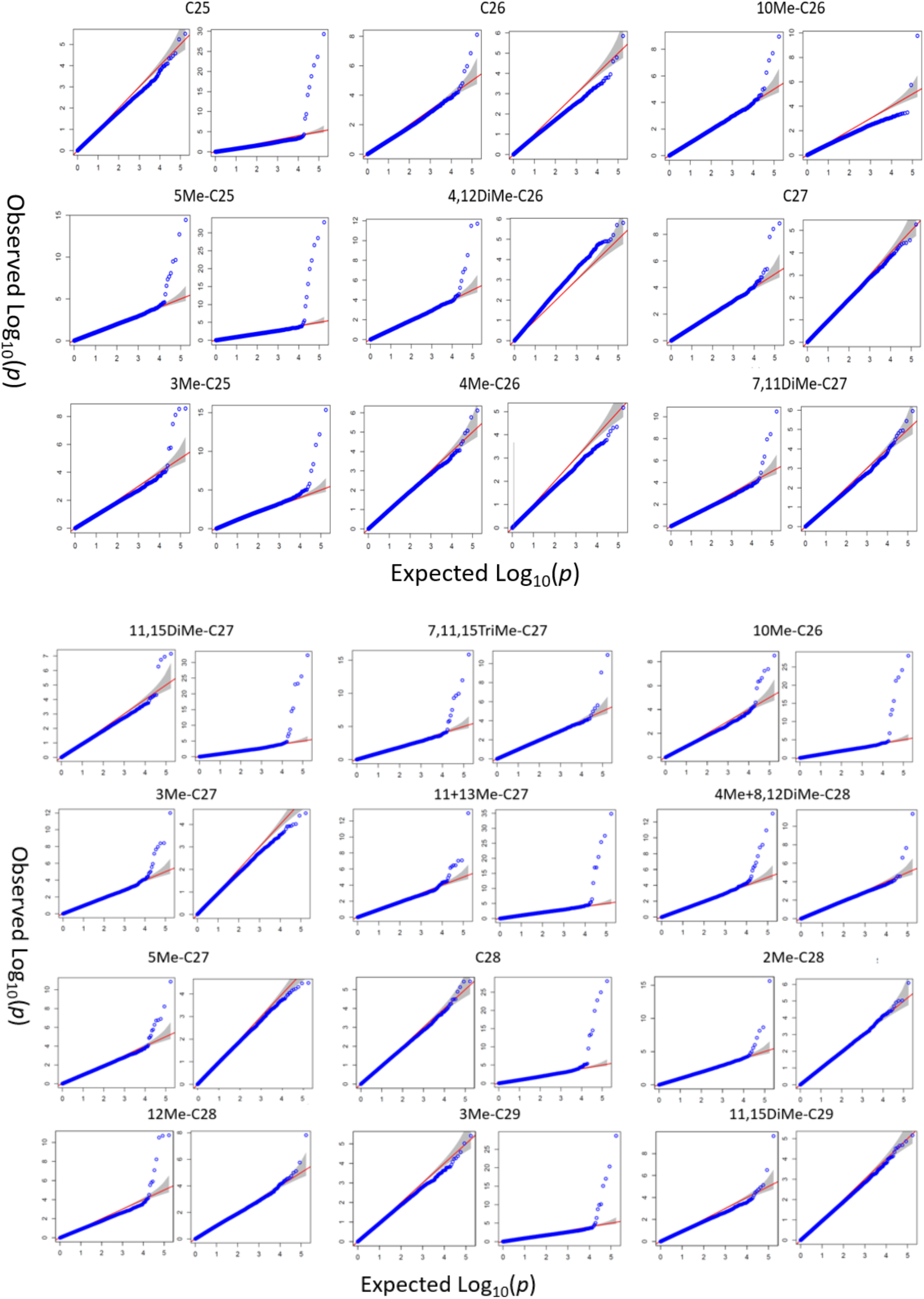
Discriminant analysis (DA) of the chemical signature of all colonies examined in this study. Samples from different nests are marked by different colors. Wilks’ Lambda test (Rao’s approximation) indicate a statistical difference between colonies (p-value<0.0001).

**Figure S3.**
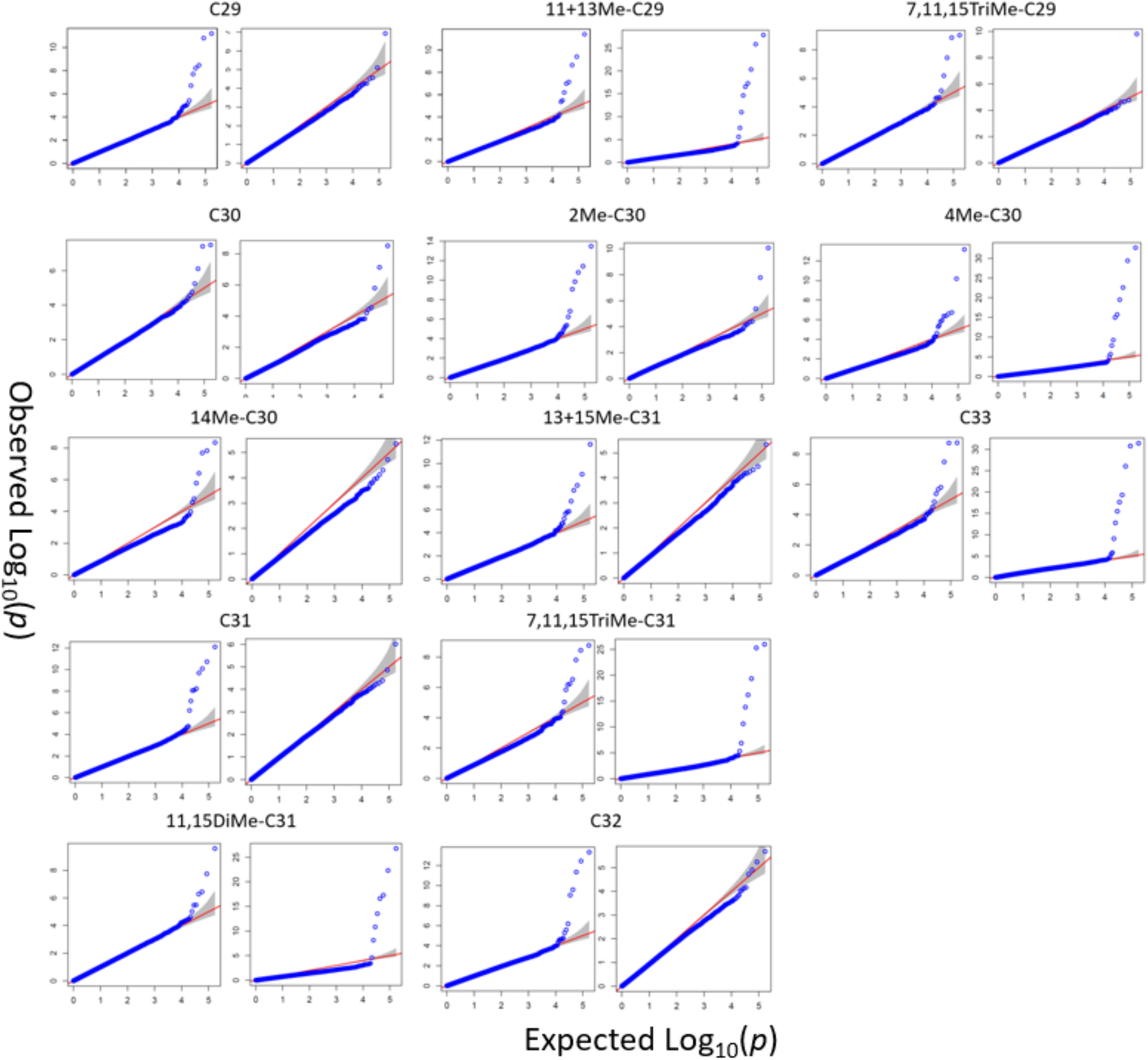
Assesment of how well the model used in the inidivual-level (left) and colony-level (right) analyses accounts for familial relatedness and population strcture for each of the CHC traits. The negative logarithms of the loci *p-*values S are plotted against their expected value under the null hypothesis of no association with the trait.

**Figure S4.**
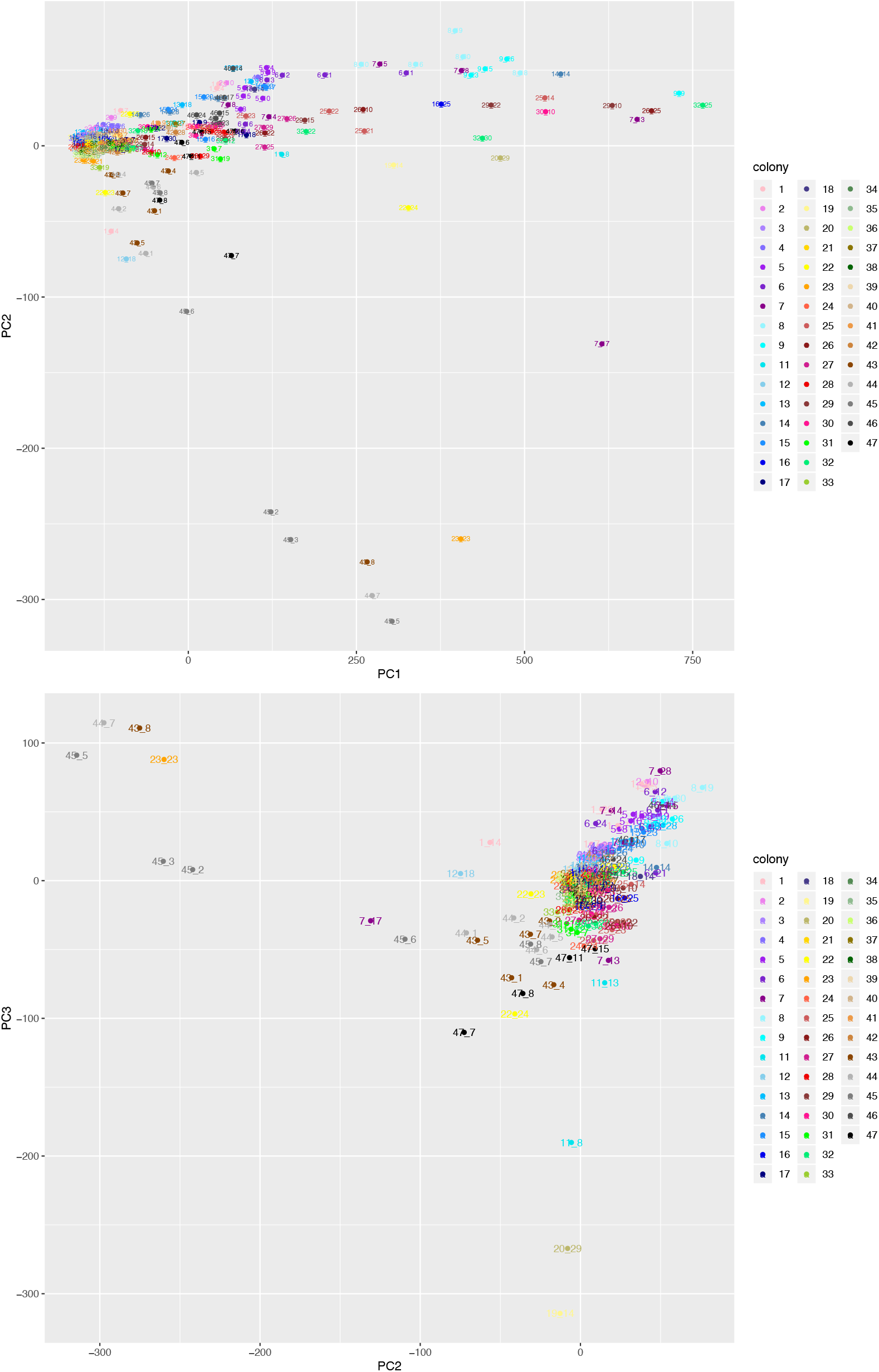
Principle Component Analysis (PCA) of genotype data from 244 individuals of 45 colonies colored by colony ID. Note that colonies were numbered from north to south along the transect.

**Figure S5.**
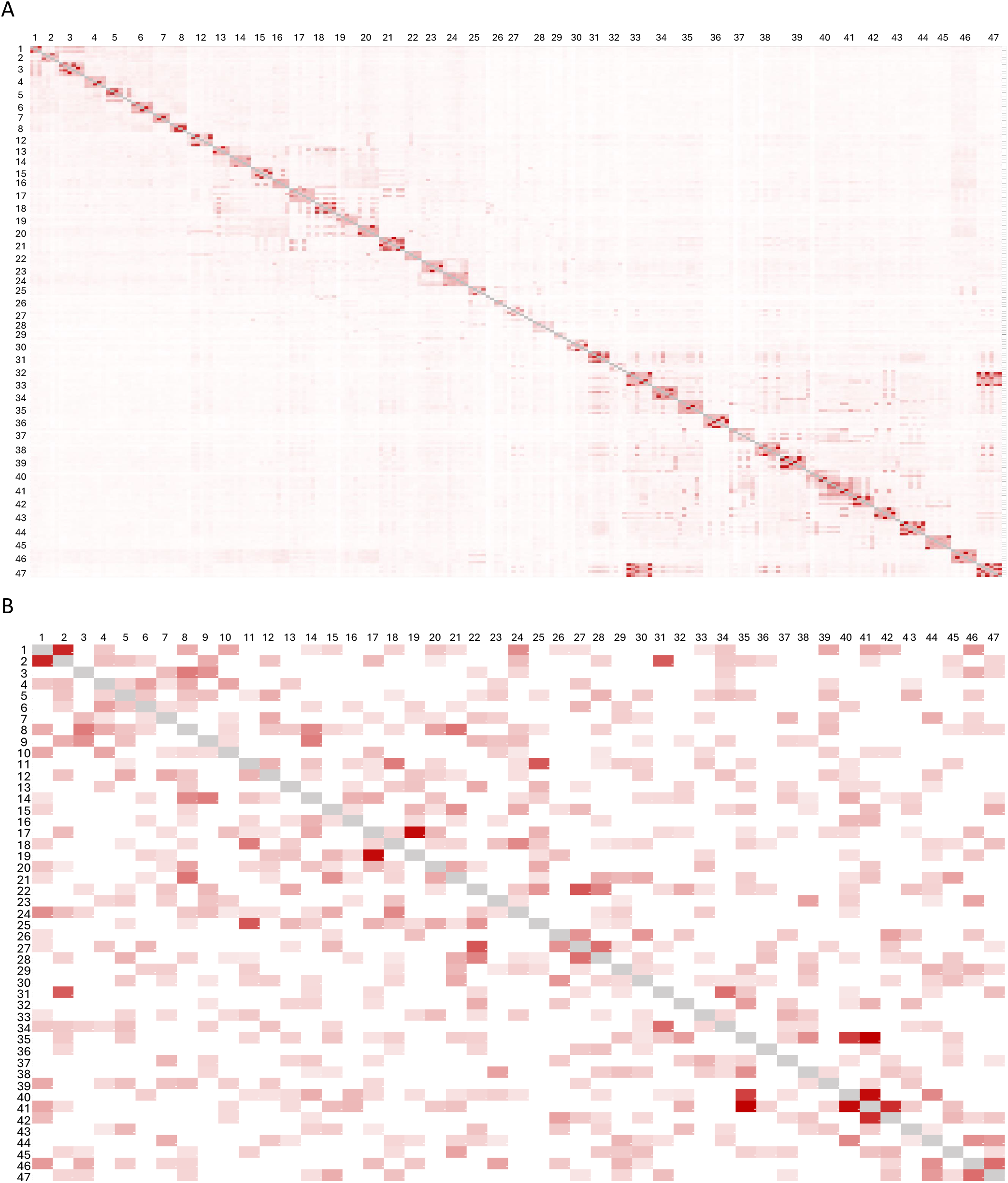
Kinship matrices based on the total length of shared IBD haplotypes between 244 individuals of 45 colonies (A) and 47 individuals of 47 colonies (B). For visibility, only colony numbers are presented. The heatmap coloring is based on the total number of bases in shared haplotypes between each pair of samples. Higher values are stronger red.

**Table S1.** Outlayer removals and normalization methods of the phenotype on the individual and colony level.

| Trait | Individual-level outlayers | Individual-level normalization | Colony-level outlayers | Colony-level normalization |
| --- | --- | --- | --- | --- |
| C25 | 17 | Yeo-Johnson | 0 | $\text{Log}_b(x + a)$ |
| 5Me-C25 | 44 | Order norm | 9 | $\text{Log}_b(x + a)$ |
| 3Me-C25 | 22 | $\text{Log}_b(x + a)$ | 4 | Box Cox |
| C26 | 23 | $\text{sqrt}(x + a)$ | 2 | Box Cox |
| 10Me-C26 | 13 | $\text{Log}_b(x + a)$ | 2 | $\text{sqrt}(x + a)$ |
| 4Me-C26 | 20 | Order norm | 3 | $\text{Log}_b(x + a)$ |
| 4,12DiMe-C26 | 12 | $\text{asinh}(x)$ | 1 | $\text{sqrt}(x + a)$ |
| C27 | 11 | Order norm | 2 | $\text{Log}_b(x + a)$ |
| 5Me-C257 | 5 | $\text{asinh}(x)$ | 0 | $\text{sqrt}(x + a)$ |
| 11+13Me-C27 | 13 | Order norm | 2 | $\text{Log}_b(x + a)$ |
| 11,15DiMe-C27 | 37 | Order norm | 9 | $\text{Log}_b(x + a)$ |
| 7,11DiMe-C27 | 2 | Yeo-Johnson | 0 | $\text{asinh}(x)$ |
| 3Me-C27 | 16 | Box Cox | 1 | Box Cox |
| 7,11,15TriMe-C27 | 21 | $\exp(x)$ | 4 | $\text{asinh}(x)$ |
| C28 | 37 | Order norm | 4 | $\text{sqrt}(x + a)$ |
| 12Me-C28 | 13 | Order norm | 0 | $\text{asinh}(x)$ |
| 4,12DiMe-C28 | 1 | Box Cox | 0 | $\text{sqrt}(x + a)$ |
| 4Me+8,12DiMe-C28 | 16 | $\text{sqrt}(x + a)$ | 0 | Box Cox |
| 2Me-C28 | 23 | Order norm | 2 | $\text{sqrt}(x + a)$ |
| C29 | 14 | Box Cox | 1 | $\text{sqrt}(x + a)$ |
| 11+13Me-C29 | 13 | Box Cox | 1 | Box Cox |
| 11,15DiMe-C29 | 17 | Yeo-Johnson | 2 | $\text{Log}_b(x + a)$ |
| 3Me-C29 | 34 | Order norm | 3 | $\text{sqrt}(x + a)$ |
| 7,11,15TriMe-C29 | 5 | Box Cox | 3 | $\text{sqrt}(x + a)$ |
| C30 | 26 | Yeo-Johnson | 4 | $\text{asinh}(x)$ |
| 14Me-C30 | 15 | Box Cox | 2 | $\text{asinh}(x)$ |
| 4Me-C30 | 16 | Order norm | 0 | $\text{Log}_b(x + a)$ |
| 2Me-C30 | 14 | $\text{asinh}(x)$ | 4 | $\text{Log}_b(x + a)$ |
| C31 | 14 | $\text{Log}_b(x + a)$ | 3 | Box Cox |
| 13+15Me-C31 | 11 | $\exp(x)$ | 4 | $\text{sqrt}(x + a)$ |
| 11,15DiMe-C31 | 11 | Box Cox | 0 | $\text{Log}_b(x + a)$ |
| 7,11,15TriMe-C31 | 4 | Box Cox | 0 | $\text{asinh}(x)$ |
| C32 | 13 | $\text{asinh}(x)$ | 4 | $\text{asinh}(x)$ |
| C33 | 8 | $\text{asinh}(x)$ | 0 | $\text{sqrt}(x + a)$ |

**Table S2.** The mapping of significant loci (*q*-value < 0.3) on the genome. Individual-level GWAS results are in blue and colony-level GWAS results are in black. Loci that are mapped to the same or close genetic position (within 12cM) are marked with yellow or orange backgrounds. Both genetic and physical positioning are shown.

| Chrom | Genetic pos (cM) | Scaffold | Position | CHC | q-value |
| --- | --- | --- | --- | --- | --- |
| 1 | 0 | scaffold17\|size760598 | 515848 | 4,12DiMe-C26 | 0.279715055 |
| 1 | 2.87 | scaffold35\|size591243 | 473891 | 4,12DiMe-C26 | 0.11352072 |
| 1 | 9.66 | scaffold296\|size210905 | 44019 | 4,12DiMe-C26 | 0.279715055 |
| 1 | 34.15 | scaffold578\|size105271 | 38187 | 7,11,15TriMe-C27 | 0.168010666 |
| 1 | 88.15 | scaffold916\|size58012 | 4774 | C31 | 0.118061549 |
| 1 | 88.15 | scaffold293\|size211391 | 198932 | C33 | 0.281531387 |
| 2 | 65.4 | scaffold636\|size94593 | 82725 | 3Me-C25 | 0.281531387 |
| 3 | 85.67 | scaffold320\|size199217 | 33276 | 11+13Me-C27 | 0.259138689 |
| 3 | 87.52 | scaffold1113\|size42492 | 15042 | 4Me+8,12DiMe-C28 | 0.168010666 |
| 4 | 1.88 | scaffold1163\|size40329 | 39019 | 4Me+8,12DiMe-C28 | 0.201612799 |
| 4 | 16.6 | scaffold558\|size111151 | 58293 | 11+13Me-C29 | 0.145481369 |
| 4 | 39.56 | scaffold77\|size466230 | 39015 | 4,12DiMe-C26 | 0.23006866 |
| 4 | 39.56 | scaffold551\|size113931 | 79453 | C31 | 0.144009142 |
| 4 | 83.57 | scaffold326\|size196711 | 73375 | 11+13Me-C27 | 0.009092586 |
| 4 | 143.72 | scaffold220\|size265453 | 70637 | C32 | 0.163469837 |
| 4 | 181.71 | scaffold734\|size79827 | 17289 | 7,11,15TriMe-C27 | 0.23793943 |
| 4 | 203.11 | scaffold74\|size472645 | 164044 | C31 | 0.118061549 |
| 5 | 131.68 | scaffold35\|size591243 | 43412 | 5Me-C25 | 0.106385132 |
| 6 | 70.38 | scaffold336\|size190304 | 25911 | C29m3 | 0.181851711 |
| 6 | 90.66 | scaffold737\|size79726 | 15861 | C28 | 0.251561534 |
| 7 | 52.75 | scaffold107\|size407829 | 186705 | C33 | 0.281531387 |
| 7 | 56.43 | scaffold859\|size63718 | 3120 | 7,11,15TriMe-C27 | 0.168010666 |
| 7 | 57.35 | scaffold19\|size722566 | 429439 | C31 | 0.144009142 |
| 7 | 62.08 | scaffold279\|size219011 | 173386 | 11,15DiMe-C27 | 0.018185171 |
| 7 | 101.52 | scaffold30\|size631625 | 144566 | C28 | 0.275375449 |
| 7 | 104.36 | scaffold85\|size450463 | 361438 | 11+13Me-C29 | 0.175789988 |
| 7 | 111.1 | scaffold28\|size641708 | 145558 | 4Me+8,12DiMe-C28 | 0.21114854 |
| 7 | 135.68 | scaffold404\|size159223 | 63505 | 3Me-C25 | 0.281531387 |
| 8 | 126.41 | scaffold194\|size285062 | 81296 | 11+13Me-C29 | 0.175789988 |
| 8 | 137.97 | scaffold618\|size98603 | 6938 | C31 | 0.118061549 |
| 8 | 208.22 | scaffold658\|size91109 | 44118 | 11,15DiMe-C27 | 0.297782177 |
| 8 | 222.43 | scaffold740\|size79674 | 67018 | C32 | 0.096871015 |
| 8 | 259.2 | scaffold20\|size721085 | 484650 | C29 | 0.28516405 |
| 8 | 309.3 | scaffold751\|size78167 | 6597 | 7,11,15TriMe-C29 | 0.287585825 |
| 9 | 45.61 | scaffold54\|size511285 | 140799 | 5Me-C25 | 0.009092586 |
| 9 | 85.74 | scaffold235\|size251712 | 52728 | C32 | 0.090816576 |
| 9 | 119.36 | scaffold219\|size265940 | 180069 | 7,11,15TriMe-C27 | 0.168010666 |
| 9 | 137.13 | scaffold311\|size203550 | 120617 | C29 | 0.186173981 |
| 9 | 142.85 | scaffold505\|size124814 | 43537 | 12Me-C28 | 0.093843796 |
| 9 | 142.85 | scaffold66\|size491290 | 166504 | 4Me+8,12DiMe-C28 | 0.276082392 |
| 10 | 105.07 | scaffold306\|size205911 | 65549 | 12Me-C28 | 0.093843796 |
| 10 | 132.88 | scaffold75\|size469975 | 364802 | 4Me+8,12DiMe-C28 | 0.21114854 |
| 11 | 2.02 | scaffold146\|size337763 | 226678 | 7,11,15TriMe-C27 | 0.271314522 |
| 11 | 17.44 | scaffold291\|size213446 | 9721 | 4Me-C30 | 0.099898234 |
| 11 | 44.83 | scaffold22\|size714790 | 90604 | 11+13Me-C27 | 0.275808429 |
| 11 | 111.65 | scaffold67\|size489733 | 28054 | C31 | 0.189579603 |
| 11 | 153.29 | scaffold622\|size97615 | 36744 | 3Me-C25 | 0.292883459 |
| 11 | 312.17 | scaffold508\|size123518 | 117137 | 12Me-C28 | 0.093843796 |
| 12 | 0 | scaffold730\|size80008 | 50311 | 2Me-C30 | 0.240663927 |
| 12 | 67.25 | scaffold71\|size482159 | 21063 | 4Me+8,12DiMe-C28 | 0.21114854 |
| 12 | 77.84 | scaffold71\|size482159 | 252805 | 4Me+8,12DiMe-C28 | 0.215941637 |
| 12 | 87.97 | scaffold593\|size103847 | 95748 | 11+13Me-C29 | 0.19614006 |
| 12 | 94.99 | scaffold65\|size491780 | 101529 | 7,11,15TriMe-C27 | 0.271314522 |
| 12 | 115.45 | scaffold209\|size272198 | 146941 | 5Me-C25 | 0.045408288 |
| 12 | 141.58 | scaffold470\|size137511 | 24197 | C31 | 0.226032368 |
| 14 | 40.37 | scaffold44\|size549762 | 519588 | 11+13Me-C29 | 0.175789988 |
| 14 | 48.21 | scaffold581\|size104792 | 64268 | 5Me-C25 | 0.207830527 |
| 14 | 50.17 | scaffold125\|size365959 | 224995 | 7,11,15TriMe-C29 | 0.254286414 |
| 14 | 92.55 | scaffold204\|size276989 | 165280 | 4Me+8,12DiMe-C28 | 0.072653261 |
| 14 | 95.38 | scaffold144\|size340398 | 186437 | 5Me-C25 | 0.141844335 |
| 14 | 107.79 | scaffold3\|size1157121 | 1010964 | 2Me-C30 | 0.265638486 |
| 14 | 127.94 | scaffold130\|size360227 | 149410 | 4Me+8,12DiMe-C28 | 0.168010666 |
| 15 | 52.68 | scaffold72\|size474807 | 247511 | 4,12DiMe-C26 | 0.11352072 |
| 15 | 90.15 | scaffold124\|size367063 | 14584 | 5Me-C25 | 0.093086991 |
| 15 | 94.21 | scaffold99\|size421651 | 410341 | 5Me-C25 | 0.027244973 |
| 15 | 174.71 | scaffold212\|size271006 | 160026 | 11+13Me-C27 | 0.172551495 |
| 16 | 40.47 | scaffold1\|size1555347 | 1092532 | 11,15DiMe-C27 | 0.297782177 |
| 17 | 55.6 | scaffold299\|size209978 | 118401 | C31 | 0.118061549 |
| 17 | 68.45 | scaffold29\|size639771 | 423039 | 5Me-C25 | 0.072740685 |
| 17 | 147.15 | scaffold608\|size100611 | 57366 | 7,11,15TriMe-C27 | 0.018163315 |
| 18 | 74.77 | scaffold153\|size330014 | 258009 | 12Me-C28 | 0.220230198 |
| 18 | 117.35 | scaffold61\|size494520 | 99800 | 11+13Me-C29 | 0.175789988 |
| 18 | 136.63 | scaffold13\|size775888 | 175137 | 5Me-C25 | 0.141844335 |
| 18 | 148.94 | scaffold74\|size472645 | 467205 | C28 | 0.227314639 |
| 20 | 0 | scaffold1423\|size26545 | 12315 | 5Me-C25 | 0.181851711 |
| 20 | 83.75 | scaffold53\|size513719 | 234805 | 4Me+8,12DiMe-C28 | 0.108979892 |
| 20 | 110.88 | scaffold392\|size162570 | 49495 | 5Me-C25 | 0.045462928 |
| 21 | 56.5 | scaffold52\|size514423 | 369128 | 7,11,15TriMe-C29 | 0.254286414 |
| 22 | 85.98 | scaffold271\|size223719 | 127531 | C28 | 0.24549981 |
| 23 | 19.07 | scaffold16\|size761867 | 381681 | 3Me-C25 | 0.281531387 |
| 23 | 45.16 | scaffold180\|size296944 | 252701 | C29 | 0.28516405 |
| 23 | 252.96 | scaffold588\|size104489 | 7055 | C29 | 0.28516405 |
| 23 | 278.7 | scaffold583\|size104635 | 55219 | C28 | 0.275375449 |
| 23 | 346.55 | scaffold27\|size651064 | 287045 | C32 | 0.174367827 |
| 23 | 347.46 | scaffold113\|size393933 | 273321 | 2Me-C30 | 0.118061549 |
| 23 | 370.92 | scaffold232\|size254158 | 2527 | 5Me-C25 | 0.106385132 |
| 24 | 143.62 | scaffold323\|size197691 | 182834 | 2Me-C30 | 0.265638486 |
| 24 | 189.98 | scaffold628\|size95885 | 66061 | C29 | 0.186173981 |
| 24 | 243.79 | scaffold215\|size268708 | 257542 | 11+13Me-C29 | 0.232997505 |
| 24 | 284.75 | scaffold26\|size661983 | 198368 | 5Me-C25 | 0.106385132 |
| 25 | 67.31 | scaffold6\|size960371 | 700278 | C32 | 0.090816576 |
| 25 | 84.71 | scaffold155\|size327501 | 273726 | C31 | 0.144009142 |
| 25 | 94.57 | scaffold19\|size722566 | 309358 | C28 | 0.261411835 |
| 26 | 21.23 | scaffold553\|size113576 | 90729 | 11,15DiMe-C27 | 0.297782177 |
| 26 | 33.21 | scaffold33\|size606826 | 197846 | 12Me-C28 | 0.292429376 |
| 26 | 127.17 | scaffold804\|size70630 | 40423 | 5Me-C25 | 0.093086991 |
| 26 | 221.23 | scaffold57\|size507111 | 381549 | C28 | 0.275375449 |
| 26 | 278.97 | scaffold303\|size207491 | 10542 | 11+13Me-C29 | 0.145481369 |
| 26 | 309.97 | scaffold8\|size894206 | 191498 | 7,11,15TriMe-C27 | 0.264881681 |

